# *In silico* identification of switching nodes in metabolic networks

**DOI:** 10.1101/2023.05.17.541195

**Authors:** Francis Mairet

## Abstract

Cells modulate their metabolism according to environmental conditions. A major challenge to better understand metabolic regulation is to identify, from the hundreds or thousands of molecules, the key metabolites where the re-orientation of fluxes occurs. Here, a method called ISIS (for *In Silico* Identification of Switches) is proposed to locate these nodes in a metabolic network, based on the analysis of a set of flux vectors (obtained e.g. by parsimonious flux balance analysis with different inputs). A metabolite is considered as a switch if the fluxes at this point are redirected in a different way when conditions change. The soundness of ISIS is shown with four case studies, using both core and genome-scale metabolic networks of *Escherichia coli, Saccharomyces cerevisiae* and the diatom *Phaeodactylum tricornutum*. Through these examples, we show that ISIS can identify hot-spots where fluxes are reoriented. Additionally, switch metabolites are deeply involved in post-translational modification of proteins, showing their importance in cellular regulation. In *P. tricornutum*, we show that Erythrose 4-phosphate is an important switch metabolite for mixotrophy suggesting the importance of this metabolite in the non-oxidative pentose phosphate pathway to orchestrate the flux variations between glycolysis, the Calvin cycle and the oxidative pentose phosphate pathway when the trophic mode changes. Finally, a comparison between ISIS and reporter metabolites identified with transcriptomic data confirms the key role of metabolites such as L-glutamate or L-aspartate in the yeast response to nitrogen input variation. Overall, ISIS opens up new possibilities for studying cellular metabolism and regulation, as well as potentially for developing metabolic engineering.

## Introduction

Despite huge developments in the last decades, deciphering cellular metabolism remains a great challenge, essential for a wide range of fields (including biotechnology, health, ecology, etc.). The analysis of metabolic networks plays an important role in addressing this issue. More and more networks are now available, with a more complete coverage of the metabolism [10]. There are also a growing number of methods for analyzing and exploiting these networks [21, 30]. Among them, one of the most common approaches is Flux Balance Analysis (FBA), which aims to predict the metabolic fluxes [23]. This method corresponds to a linear programming problem, where one wants to maximize or minimize an objective function such as biomass synthesis or ATP production, given steady-state conditions and a set of flux constraints on e.g. substrate uptake.

Although FBA and all its derivatives have been successfully used, the size of the networks (now often with several thousands metabolites and reactions) makes their analysis or even the exploitation and visualization of the results more and more complex [16, 6]. To help explore these networks, the identification of key metabolites is a major concern [31]. Here, we will focus on the nodes in the networks where the fluxes are redirected when culture conditions vary. Actually, the cellular response to environmental changes such as nutrient inputs leads to a reorganization of metabolism, which can be difficult to map given the scale of the network. By highlighting where the main changes in metabolism occur, the identified switching nodes will give us a better picture of cellular response to environmental variations and could help in the modelling, monitoring or control of cell systems.

The identification of key metabolites has been the subject of many studies (e.g. [19, 14, 27, 17]), although there is no general agreement about the definition of what a key metabolite is. Following approaches developped for network analysis, key metabolites can be defined by their degree, i.e. the number of reactions in which they are involved [31]. Other aspect of the network topology can also be used, such as node centrality [19]. On the other hand, [14, 17] have focused on essentiality: a metabolite is essential if biomass cannot be produced if this metabolite is removed from the metabolic network. More in line with my vision of switch points, i.e. a point where the fluxes are reoriented in response to environmental changes, [24] provide a useful method to identify the so-called reporter metabolites. They correspond to nodes around which the enzymes are subject to the most significant transcriptional changes. Nonetheless, reporter metabolites are identified from experimental data and a purely theoretical method (based only on the metabolic network) is still lacking.

Here, I propose to identify switch nodes based on the analysis of a set of flux solutions under different conditions, obtained e.g. by parsimonious flux balance analysis (pFBA) [18]. The key idea is to determine mathematically the metabolites around which the most significant metabolic rewirings occur. To do so, for each metabolite, we consider the flux vectors (including stoichiometry) of the reactions involving this metabolite (as a substrate or a product) for all the conditions. The metabolite is considered as a switch node if the dimension of the vector space generated by this set is greater than one, i.e. if the flux vectors involving this metabolite for different conditions are not co-linear, reflecting a reorientation of the fluxes in response to changing conditions. After explaining in details how the switches are identified, the core metabolic network of *Escherichia coli* will be used to illustrate the method. Then, we will show that ISIS brings out metabolites involved in post-translational modification (PTM) of proteins in *E. coli*. ISIS will also be used to decipher how metabolism is impacted by trophic modes in the diatom *Phaeodactylum tricornutum*, and by nitrogen limitation for *Saccharomyces cerevisiae*, in comparison with the reporter metabolite method. Finally, we will assess the robustness of ISIS with respect to flux sampling.

## Results

### ISIS principle

Our objective is to identify switch nodes in a metabolic network, corresponding to key metabolites where flux reorientations occur when environmental conditions change. ISIS is based on the analysis of a set of flux vectors for a range of environmental conditions (e.g. different inputs, different objective functions reflecting different metabolic stages, etc.). A metabolite is considered as a switch if the fluxes at this point are redirected in a different way when conditions change. This is illustrated in Figure 1. On the top, the fluxes around metabolite *x*_1_ are distributed always in the same way (i.e. one third of the incoming flux *v*_1_ goes to *v*_2_, the remaining goes to *v*_3_), so *x*_1_ is not a switch node. On the contrary, in the bottom example, the incoming flux is rerouted according to conditions, so *x*_1_ is considered in this case as a switch point. This simple principle can be evaluated numerically using linear algebra (see Section Method), by evaluating the dimension of the vector space generated by the set of reaction flux vectors. More precisely, a singular value decomposition is carried out and a score between zero and one is computed from the singular values. A score of zero means that the vectors are collinear (Fig. 1, on top): the fluxes are always distributed in the same way. This already concerns all the metabolites involved in only two reactions. The higher the score, the more the metabolite can be considered a switch. All metabolites are then ranked according to their score, to highlight the most significant ones on a case-by-case basis.

**Figure 1.**
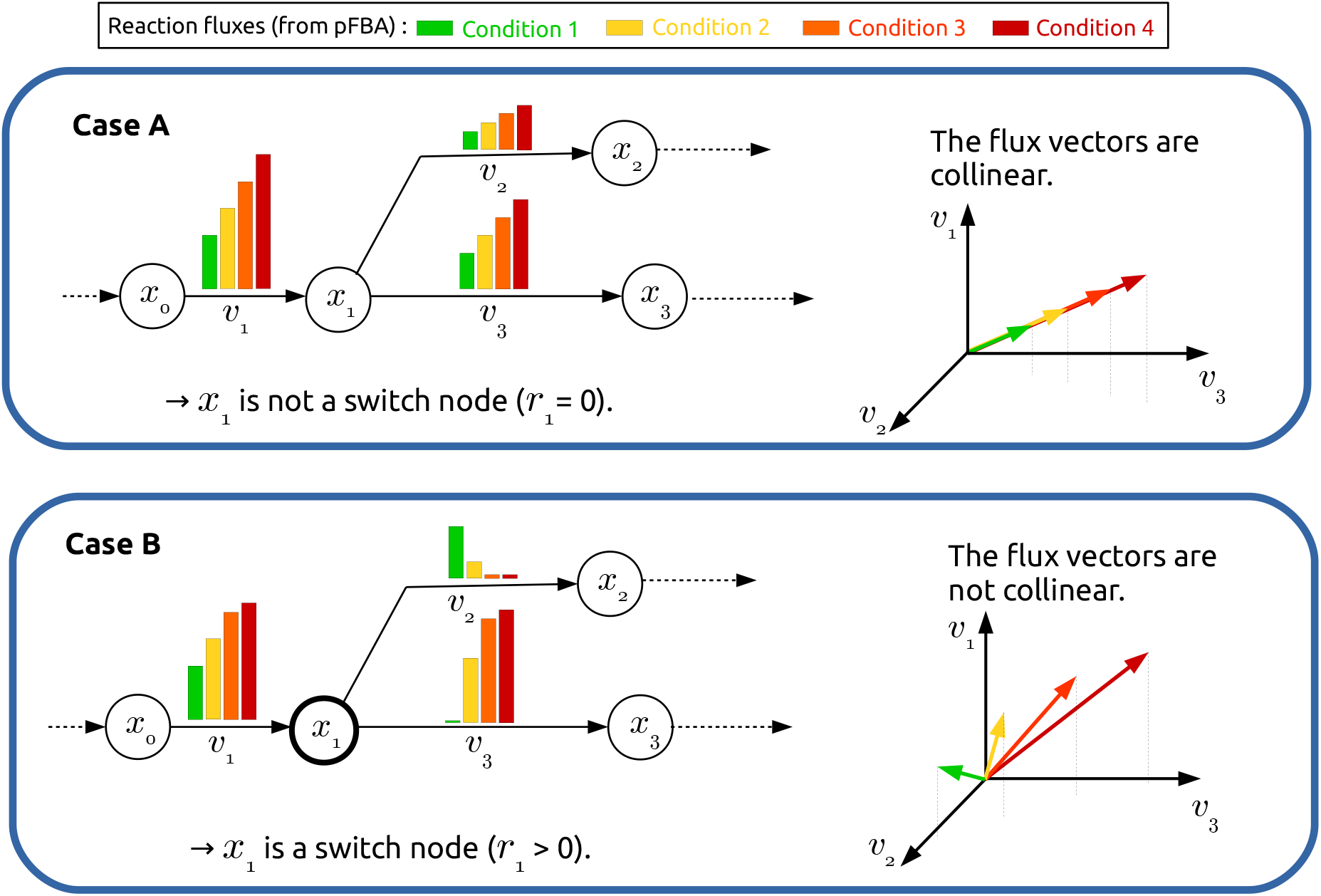
Principle for switch node identification in metabolic networks. The circles *x*_*i*_ and the arrows *v*_*j*_ represent respectively metabolites and reactions. The colored bars show reaction fluxes (computed e.g. by pFBA) for different conditions (each color represents a condition). Case A: the redistribution of fluxes around metabolite *x*_1_ occurs always in the same way, i.e. the flux vectors are collinear, so *x*_1_ is not a switch point. Case B: the incoming flux is rerouted according to conditions, so *x*_1_ is a switch point.

### A toy example: *E. coli* under aerobic vs anaerobic conditions

The principle of ISIS is first illustrated by studying the transition from aerobic to anaerobic conditions in *E. coli*. We use its core metabolic model, composed of 72 metabolites and 95 reactions [22], and estimate flux vectors with and without oxygen using pFBA (see Fig. 2). The switching scores of all the metabolites are given in SI1. ISIS has identified as switch nodes the junctions of glycolysis with the TCA cycle (pyruvate and acetyl-coA) and with the oxydative pentose phosphate pathway (glucose 6-phosphate). This is in line with what we could expect given that these last two pathways are shut down in absence of oxygen [28, 22]. We also observed that several currency metabolites (e.g. ATP, NADPH), involved in many reactions, are also identified as switch points. Most of these metabolites have already been recognized on the basis of their degree [19] or centrality [31]. On the other hand, some metabolites with a high degree or centrality such as glutamate [31] have a low score with ISIS when focusing on aerobic vs anaerobic growth. This simple example shows the soundness and capacity of ISIS to identify nodes around which the metabolism is rerouted taking into account the specific conditions laid down.

**Figure 2.**
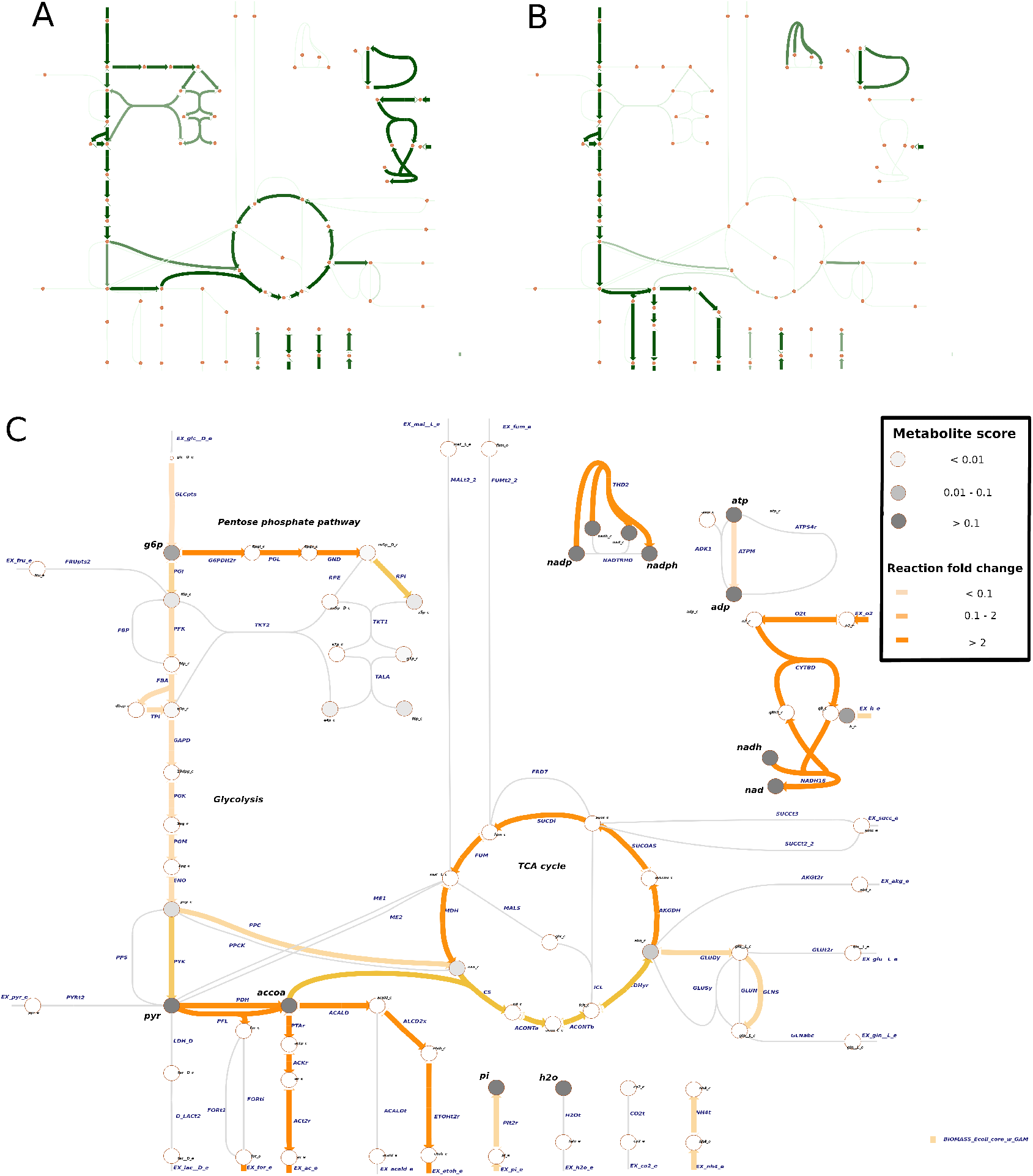
Switch nodes for *E. coli* under aerobic vs anaerobic conditions. Overview of metabolic fluxes under aerobic (A) and anaerobic (B) condition (see C for the name of metabolites and reactions). C: Comparison between these two conditions. Switch nodes (dark dots) have been identified at the junctions of glycolysis with the TCA cycle (pyruvate and acetyl-coA) and with pentose phosphate pathway (glucose 6-phosphate).

### ISIS brings out metabolites involved in protein PTM in *E. coli*

Given that the switch metabolites are by definition nodes where fluxes are reoriented, these metabolites can potentially be involved in cellular regulations, such as PTM. To investigate this aspect, we follow [4], by considering the growth of *E. coli* under 174 different nutrient inputs. Using a network composed of 2382 reactions and 1668 metabolites [8], we first define the flux vectors for each input using pFBA. We then analyze all these fluxes with ISIS. The list of the switching points is given in SI2. Among the top score, we find key branching points between the main pathways, such as Fructose 6-Phosphate, Pyruvate, Acetyl-CoA, etc.

On Fig. 3, we plot the proportion of metabolites known to be ligands of protein (i.e. involved in PTM), according to their switching score. Ligands are clearly enriched among the top switching metabolites (hypergeometric test, p=6.10^−8^). For the class of metabolites with the highest scores (>0.3), 74% of the metabolites are ligands, compared to 25% for the class with the lowest scores (*<*0.1).

**Figure 3.**
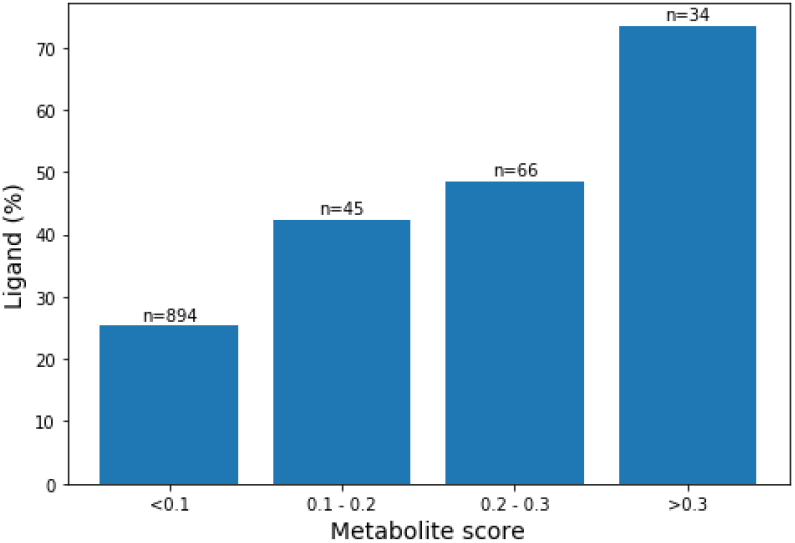
Percentage of metabolites identified as ligand in protein PTM as a function of their scores in ISIS for *E. coli*, based on the fluxes for 174 different inputs with the genome-scale metabolic network iAF1260 [8]. Metabolites identified as switch point are much more involved in PTM, highlighting their role in cellular regulation.

**Figure 4.**
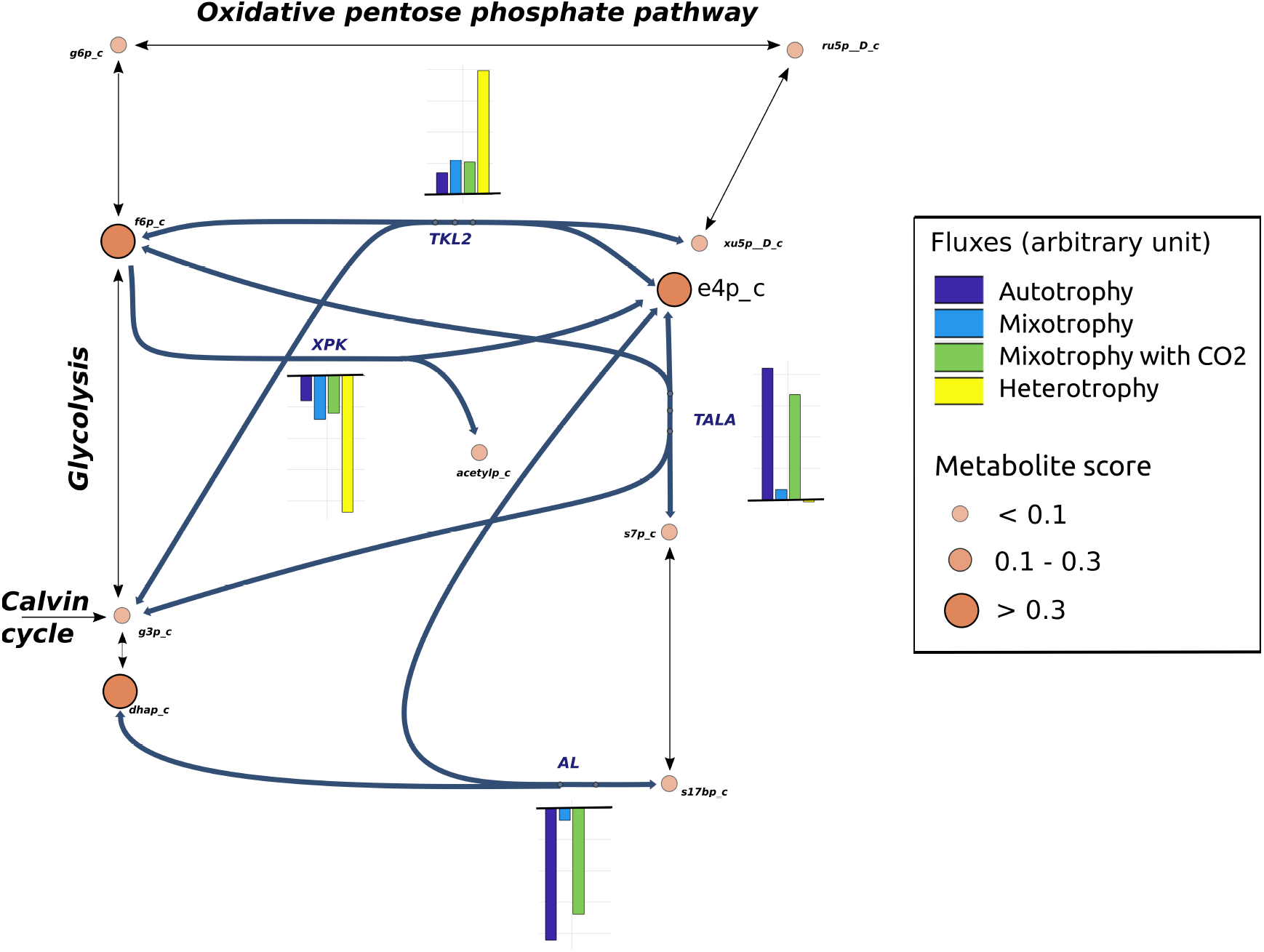
Metabolic fluxes in the non-oxidative pentose phosphate pathway for different trophic modes in the diatom *P. tricornutum*. Alongside dihydroxyacetone phosphate (dhap) and fructose-6-phosphate (f6p), erythrose 4-phosphate (e4p) appears as a key hub to balance the fluxes between glycolysis, the Calvin cycle and the oxidative pentose phosphate pathway, showing the role of the non-oxidative pentose phosphate pathway in orchestrating the energy balance of the cell.

Based on a comparison of FBA fluxes, [4] have identified highly regulated reactions, which appear to account for a significant proportion of the known proteins with PTM. By contrast, here we clearly show the key role of the switching metabolites identified by ISIS in protein PTM. Overall, the identification of switch nodes could help in deciphering the complex roles of metabolites in the regulation of protein activity [32].

### Shedding light on mixotrophy in the diatom *P. tricornutum*

We study the effect of trophic mode in *P. tricornutum*. The fluxes for autotrophic, mixotrophic and heterotrophic^1^ growths have been estimated in [13], considering a genome-scale metabolic network composed of 587 metabolites and 849 reactions. This study highlighted the importance of flux rerouting between chloroplasts and mitochondria, depending on the trophic mode. Using this set of fluxes, ISIS identifies as switch nodes glycerone-P (dihydroxyacetone phosphate), fructose-6-phosphate, fructose 1,6-bisphosphate, fumarate, pyruvate, etc. (see SI 3). These metabolites are key branching points between chloroplast and mitochondria (as shown in Fig. 1 from [13]). Additionally, we also identify erythrose 4-phosphate, which is actually a major hub of the metabolism to balance the fluxes between the main pathways (in particular glycolysis, the Calvin cycle and the oxidative pentose phosphate pathway). Thus, ISIS contributes to our understanding of trophic modes by suggesting the overlooked role of a key metabolite in the non-oxidative pentose phosphate pathway in orchestrating changes between the energetic pathways.

### ISIS identifies reporter metabolites for *Saccharomyces cerevisiae* under nitrogen limitation

Given that our definition of switch nodes is close to reporter metabolites [24], we compare both methods, using [29] as a case study. To do so, we consider the growth of *S. cerevisiae* under three different nitrogen limitation: ammonium 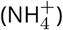, alanine (Ala), and glutamine (Gln). For each input, the flux vectors are computed with pFBA using the metabolic network YeastGEM v8.1.1 composed of 2241 metabolites and 3520 reactions [12]. ISIS is then applied by paire-wise comparison (Gln vs 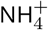, Ala vs 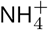, and Ala vs Gln), following what have been done in [29]. The score of all the metabolites for the three comparisons are given in SI 4. Switch metabolites with a score above 0.1, corresponding to between 11 and 16 metabolites out of 2241, are shown in Fig. 5 together with the ten first reporter metabolites given in [29]. Many of the switch metabolites, common between the different cases, are involved in amino acid synthesis. We also observe that reporter metabolites are enriched among the top switch metabolites (hypergeometric test, p=0.048, 0.061, and 3.10^−5^ for respectively Gln vs 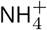, Ala vs 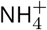, and Ala vs Gln). When comparing Ala versus Gln limitations, glutamate, aspartate and 2-oxoglutarate are depicted with both methods. The first two metabolites are closely related to the sources of nitrogen. The last one points out the importance of the TCA cycle in the synthesis of amino acids, as reported in [29]. Finally, several reporter metabolites - in particular those highlighted in [29] - appears in the top of the switch metabolite list (see SI 4), even if they do not appear in Fig. 5. For example, in the comparison between Ala versus 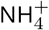, the reporter metabolites pyruvate and Acetyl-CoA are ranked respectively 19 and 46 over 2241 metabolites by ISIS. On the other hand, some reporter metabolites are not identified as switch, pointing out some differences between the two approaches. As a striking example, allantoin was reported as the reporter metabolite with the highest score in the Ala vs 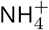 comparison, without however figuring out why it emerges. This metabolite is involved in only two reactions in our metabolic network, so it cannot be identified as a metabolic switch by ISIS. Other reporter metabolites present a low switch score, such as those involved in lipid metabolism in the Gln vs NH4 comparison. This suggests that the nitrogen input change leads to a modification of lipid metabolism that pFBA does not predict.

**Figure 5.**
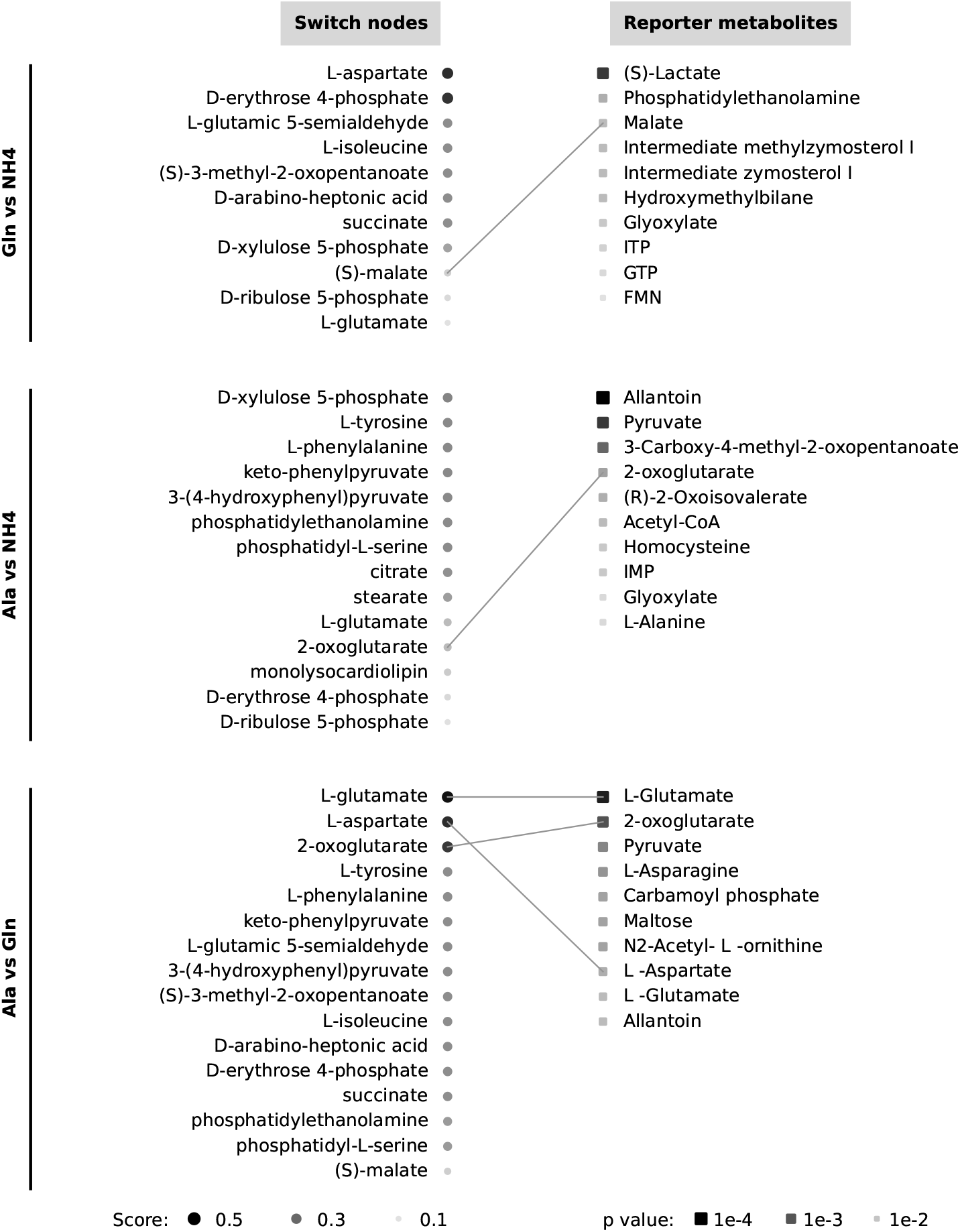
The main switch nodes identified by ISIS for *Saccharomyces cerevisiae* under nitrogen limitation, in comparison with reporter metabolites computed from transcriptomic data [29].

To conclude, this case study shows that some key metabolites can be identified *in silico*, without requiring experiments. ISIS can also complement the reporter metabolite approach, which requires transcriptomic data. Comparison between the two methods can confirm the role of a number of key metabolites and identify doubts about some of them, or more indirect effects that are difficult to predict.

### Robustness to flux sampling

The identification of switch points is based on a set of flux vectors obtained in silico. Nonetheless, different solutions can give the same objective, even with pFBA. In the following, we explore how this variability can impact the identification of switch points. Instead of taking one solution per condition, we sample the flux spaces and compute switch scores for several combinations of flux vectors chosen randomly. We test this approach with two previous case studies: the core metabolic network of *E. coli* with or without oxygen and the genome-scale network of *S. cerevisiae* with different nitrogen sources. In the first case, the switch scores obtained with flux sampling are very similar to those with pFBA (see Fig. 6 ans SI 5). Nonetheless, they show a high standard deviation, reflecting the effect of flux sampling, although this does not affect switch identification. Given the small size of this network, the flux variability is limited so the switch points remain the same. For the genome-scale network of *S. cerevisiae*, the results are more different. Figure 6 shows the comparison for Ala vs Gln. The other cases are given in SI 7 and 8, as well as the list of switch metabolites in SI 6. On one hand, most of the switch points identified previously have still a high score (e.g. L-glutamate and 2-oxoglutarate for Ala vs Gln), pointing out their key role in redistributing fluxes. On the other hand, several metabolites that were not considered as switch points present a high score when considering flux sampling. However, it is necessary to assess the relevance of these new candidates. Large-scale network present several variants of the same pathway, which creates multiple switches if for each condition a different variant is used for the same pathway. For example, in lipid synthesis, many reactions can be interchanged, including alterations of compartments. This results in the emergence of numerous lipid intermediates with high switching scores that are not relevant for the change of condition under investigation (see Fig. SI 9). Exploring the flux space is therefore useful to confirm the identification of switch points, but it also leads to switch point candidates that warrant further examination.

**Figure 6.**
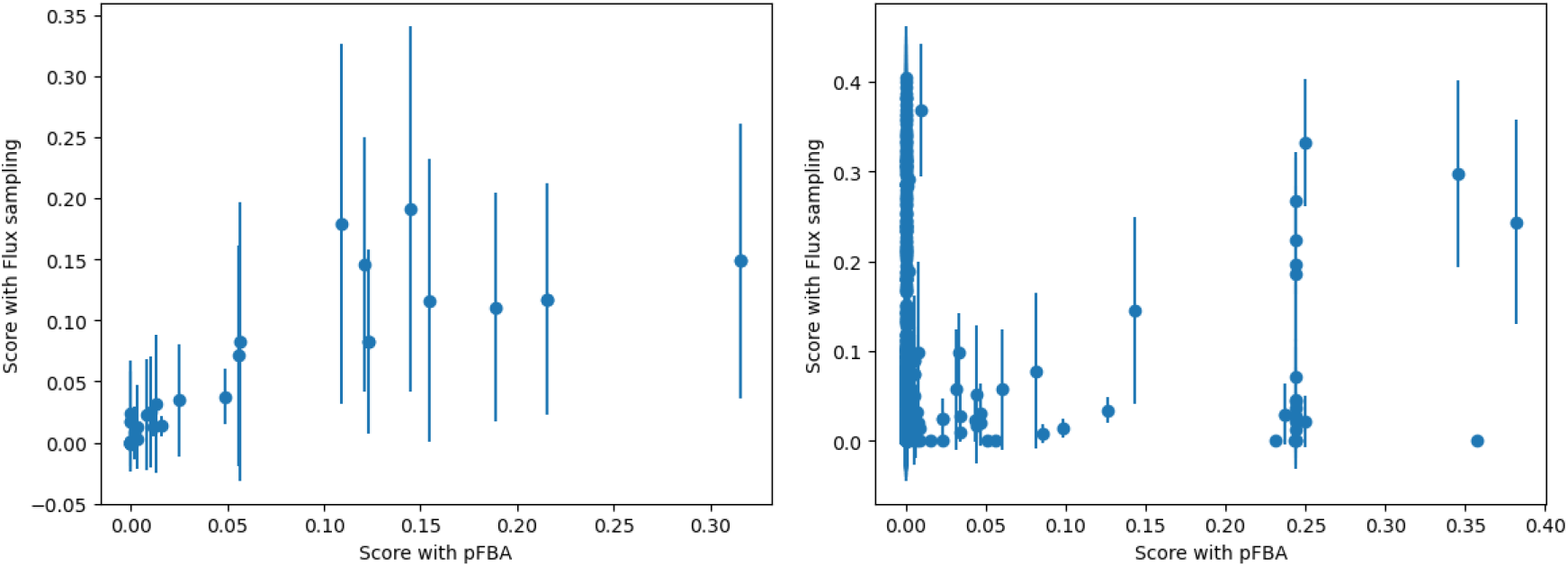
Comparison of the switch scores obtained with pFBA (on the x-axis) versus flux sampling (on the y-axis, with error bars showing standard deviation). A significant correlation was observed for the core metabolic network of *E. coli* with or without oxygen (left, Spearman correlation: *ρ* = 0.82, *p* = 5.10^−19^), but not for the genome scale network of *S. cerevisiae* with different nitrogen sources (right, for Ala vs Gln, Spearman correlation: *ρ* = 0.26, *p* = 3.10^−43^).

## Discussions

A method - called ISIS - has been proposed to identify switch nodes, i.e. metabolites around which fluxes are rerouted when environmental conditions change. These points are determined *in silico* using linear algebra: the reaction fluxes involving the metabolite under different conditions generate a vector space whose dimension reflects whether a reorientation of fluxes occurs at this node. The method is fast and scalable, e.g. it takes just a few seconds with a standard computer for a metabolic network of a few thousand reactions. ISIS gives sound results on the different case studies developed in this article. As a proof of principles, by comparing aerobic and anaerobic conditions in *E. coli*, ISIS has identified the metabolites at the junction of the main pathways, including pyruvate as one can expect [28]. A further analysis for the same species with 174 different substrates revealed that metabolites with the highest switching score are particularly involved in cellular regulation via PTM. Another important finding concerns the diatom *P. tricornutum*. When considering different trophic modes, metabolic modifications occur at different nodes to maintain the energetic balance, in particular in the non-oxydative pentose phosphate pathway with Erythrose 4-phosphate.

The analysis of metabolic networks has been the subject of several developments. Among them, a method which, like ISIS, analyses the dependencies between the fluxes is flux coupling analysis [5]. Two reactions are fully coupled if a flux for one of these reactions implies a fixed flux for the other, and vice versa. If all the reactions in which a metabolite is involved belong to a fully coupled set, that metabolite is not a switch. Flux coupling analysis can therefore be used to eliminate candidate metabolites, but does not provide a sufficient condition for identifying switches. Several methods dealing with the identification of key metabolites have also been proposed (e.g. [19, 14, 27, 17]). A comparison between them is not straightforward, given that the definition of key metabolites is not necessarily the same between the different studies. One of the closer definition is that of reporter metabolites [24]. The results on a case study with *S. cerevisiae* have shown similarities, confirming the role of some metabolites such as L-glutamate or L-aspartate as switch in response to different nitrogen inputs. Nonetheless, other reporter metabolites have low switching score. Metabolic simulations predict that no reorientation of fluxes occur at these nodes, while changes of gene expression are observed around it. This calls for future work to clarify the discrepancy on the role of these metabolites and to ascertain whether metabolic alterations occur at these points.

The main advantage of ISIS, in particular in comparison with the reporter metabolite approach, is that it does not require experimental data. The downside is that it entirely relies on flux estimations, which suffers from uncertainties at different levels (in the network reconstruction, in the definition of the biomass or the objective function, in the computation of fluxes…) [3]. A first step to overcome this limitation is to use ISIS with flux sampling, as illustrated here with *E. coli* under aerobic vs anaerobic conditions and *S. cerevisiae* under nitrogen limitations. This approach could confirm the role of some switch metabolites, but new candidates can also appear, particularly in large-scale networks, with possibly some false positives. This points out that the flux quality can strongly affect overall results. These estimations can benefit from all the recent progresses in metabolic network reconstruction and constraint-based modeling [7], in particular with the development of enzyme constraint or resource allocation methods [9, 33]. Additionally, the soundness of flux estimations can potentially be increased by integrating experimental data (e.g. transcriptomic, proteomic, fluxomic) [25], although losing the ease of a purely *in silico* approach.

Given the crucial role of switch metabolites in cellular metabolism, several applications can be considered. First, ISIS could be used to select key metabolites to monitor when studying the response of an organism to a change of environment. In the same vein, theses switch nodes can also be useful to study cellular regulations, as already illustrated by the high proportion of switch metabolites involved in PTM. The identification of switch points would also be of great interest in biotechnology: they correspond to potential targets to reorient the metabolism of the cell for a given purpose (such as the production of a metabolite of interest), either by controlling environmental conditions or by genetic manipulations. Finally, ISIS could also be used to decompose the whole network into different modules connecting the switch nodes, in order to analyze the metabolism or to develop dynamical model as proposed in [2, 1]. This would give a new approach for network splitting, complementing a set of methods (reviewed in [26]) based on network topology, flux coupling, or elementary flux mode. A specificity of our approach in that case is that the set of selected conditions defines the node identification. Thus, the same metabolic network can be decomposed in different ways depending on which conditions are considered.

To conclude, the metabolites around which fluxes are switched in response to environmental changes are key points in the metabolic network. By identifying them *in silico*, ISIS allows a better comprehension of cellular metabolism and regulation, as highlighted in this article with the studies of *E. coli, S. cerevisiae*, and *P. tricornutum* under different substrate inputs or trophic modes. This method can be easily applied to many organisms, as it only requires their metabolic network. It offers several perspectives, from fundamental studies on metabolism to biotechnology applications through metabolic engineering.

## Method

### Framework

The metabolism of a cell can be represented by its metabolic network, composed of *n*_*m*_ metabolites and *n*_*v*_ reactions. It is generally described by a stoichiometric matrix *S*(*n*_*m*_ × *n*_*v*_), where each row corresponds to a metabolite and each column to a reaction. The metabolic fluxes through this network are given by the reaction rate vector 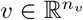. To estimate these fluxes, FBA makes two main assumptions [23]. First, the metabolism is considered at steady-state (corresponding to balanced growth condition):

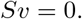

Additionally, some bounds on the metabolic fluxes are defined:

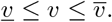

These bounds are used in particular to define nutrient inputs and to specify the reversibility of each reaction. Finally, an objective function to be maximized or minimized is considered (e.g. biomass synthesis or ATP production), defined by an objective vector 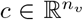. The metabolic fluxes are then the solution of a linear optimization problem (also called LP problem, for Linear Programming), which can easily be solved numerically:

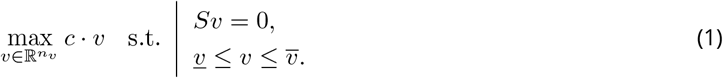

One limitation of FBA is that it can have several solutions (with the same objective value). To tackle this problem, [18] have proposed pFBA, which consists in two steps. First, FBA is used to find the optimal value of the objective function. Then, we determine the solution with the same objective value that minimizes the sum of fluxes. This tends to reduce the solution space, and it had been shown that it is consistent with gene expression measurements [18].

### ISIS principle

Switch nodes will be identified based on the analysis of a set of reaction fluxes under *n*_*c*_ different conditions, e.g. different nutrient inputs. For each condition *j* ∈ {1, …, *n*_*c*_}, the cellular metabolism is characterized by the flux vector *v*^*j*^ obtained by pFBA (or another method). Each vector is normalized, by dividing by its quadratic norm, to give each condition the same weight in the analysis.

Then, for each metabolite *i* and each condition *j*, we consider the flux vector (including stoichiometry) of the reactions involving this metabolite (as a substrate or a product) in this condition, i.e.:

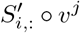

where ◯ stands for the Hadamard product (i.e. element-wise). Now given all the conditions, we consider the vector space *M*_*i*_(*n*_*v*_ × *n*_*c*_) generated by all the flux vectors for metabolite *i*:

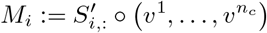

If the dimension of this vector space is greater than one, i.e. if the flux vectors involving this metabolite for different conditions are not collinear, the metabolite is considered as a switch node (see Fig. 1).

From a practical point of view, the dimension of *M*_*i*_ is evaluated by singular value decomposition (SVD), so we get:

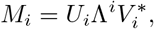

where *U*_*i*_(*n*_*v*_ × *n*_*v*_) and *V*_*i*_(*n*_*c*_ × *n*_*c*_) are unitary matrices and Λ^*i*^(*n*_*v*_ × *n*_*c*_) is a diagonal matrix whose diagonal entries 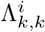 correspond to the singular values of *M*_*i*_ (ordered in descending order). We compute a score which represents the significance of each switch node:

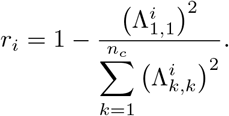

If all the vectors are collinear, then all the 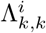 for *k >* 1 are almost null, so *r*_*i*_ ≃ 0. The closer the score is to one, the more the metabolite corresponds to a switch. The metabolites with the highest scores are finally selected.

### ISIS implementation

ISIS has been implemented under Python 3.10 within the COBRApy framework [11]. The vector fluxes for all the conditions are computed with pFBA using as objective the maximization of biomass production. A loop on all the metabolites is carried out, running for each metabolite the SVD and then computing the score. To speed up the program, all the zero rows in *M*_*i*_ are removed before running the SVD. Finally, all the metabolites are ordered following their scores. A threshold can be set above which the metabolite is considered a switch. This threshold must be adapted to each case study, in particular according to the number of conditions considered which will affect the dimension of *M*_*i*_ and therefore the overall scores.

### Case studies

ISIS has been carried out on four examples. Unless otherwise stated, the procedure described above was applied. Some specificities for each case study are given below.

#### *E. coli* (core metabolic model)

As a first case study, we consider the core metabolic network of *E. coli* [22]. We simulate growth on glucose in aerobic and anaerobic conditions (i.e. with or without oxygen). Figure 2, which compares the fluxes between the two conditions and highlights the switch metabolites, was drawn with Escher [15].

#### *E. coli* (genome-scale model)

*E. coli* network iAF1260 [8] has been used with 174 different inputs, as done in [4]. ISIS was applied by comparing all the conditions at the same time, i.e. *n*_*c*_ = 174. The objective was then to evaluate the role of switch metabolites in cellular regulation. If a metabolite is present in several compartments, its highest score is considered. We then checked the presence of the metabolites in the list of all the metabolites involved in PTM, taken from [4] (Dataset S2).

#### P. tricornutum

The metabolic network of *P. tricornutum* was taken from [13]. Four different trophic modes were considered: autotrophy, mixotrophy with or without inorganic carbon, and heterotrophy. The flux vectors, computed by FBA coupled with Euclidean norm minimization, were taken from the same article (Table S1).

#### S. cerevisiae

The metabolic network YeastGEM v8.1.1 [12] has been used, simulating aerobic growth on glucose with three different nitrogen sources: ammonium, alanine, and glutamine. This mimicks the experiments carried out in [29]. The switch metabolites are computed by comparing the nitrogen sources two by two (i.e. *n*_*c*_ = 2), to follow the aforementioned article. All the currency metabolites (ATP, NADPH, etc.) were removed to plot Fig. 5, as they are not taken into account in [29]. For each case, the results are compared with the ten reporter metabolites given in [29].

### Robustness to flux sampling

Instead of using just one vector flux, we sample the flux space for each condition using *optGpSampler* algorithm [20] from COBRApy, leading to *n*_*s*_ flux vectors per condition (we use typically *n*_*s*_ = 10^4^). We randomly define *n*_*s*_ combinations, taking one solution from the samples for each condition, and calculate the switching scores of all metabolites for each combination. Finally, we calculate for each metabolite the mean and standard deviation of the score over all the combinations. The robustness analysis has been carried out for two of the case studies: the *E. coli* core network and the *S. cerevisiae* genome-scale network.

### Statistics

The hypergeometric test was used to test for over-representation of successes, i.e. ligand for *E. coli* and reporter metabolites for *S. cerevisiae*, in the list of identified switch metabolites. The hypergeometric p-value is computed as the probability of randomly drawing k or more successes in n total draws (i.e. the number of switch metabolites), from a population of N metabolites containing K successes.

Spearman rank correlation coefficient (*ρ*) was computed to assess the relationship between the switch scores obtained with pFBA versus those obtained with flux sampling.

## Acknowledgements

The author thanks all those who provided fruitful comments during presentation of this work at JOBIM, FOSBE, and *Algae in silico* meeting, as well as the reviewers.

## Data, script, code, and supplementary information availability

All the codes (in Jupyter notebooks), data and SI are available at https://github.com/fmairet/ISIS.

- SI1: List of switch metabolites for *E. coli* under aerobic vs anaerobic conditions.
- SI2: List of switch metabolites for *E. coli* under 174 diferent nutrient inputs.
- SI3: List of switch metabolites for *P. tricornutum* under different trophic modes.
- SI4: List of switch metabolites for *S. cerevisiae* under nitrogen limitation.
- SI5: List of switch metabolites for *E. coli* with flux sampling.
- SI6: List of switch metabolites for *S. cerevisiae* with flux sampling.
- SI7: Figure. Comparison of the switch scores obtained with pFBA versus flux sampling for *S. cerevisiae* under Gln vs 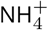.
- SI8: Figure. Comparison of the switch scores obtained with pFBA versus flux sampling for *S. cerevisiae* under Ala vs 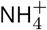.
- SI9: Figure. Alternative pathway can create false positive switch when using flux sampling, as illustrated here with an example concerning the study of *S. cerevisiae* under nitrogen limitation.

## Footnotes

1 mimicking night-time metabolism, as *P. tricornutum* do not grow normally on glucose.

